# Receptor- ligand interactions in plant inmate immunity revealed by AlphaFold protein structure prediction

**DOI:** 10.1101/2024.06.12.598632

**Authors:** Li Wang, Yulin Jia, Aron Osakina, Kenneth M. Olsen, Yixiao Huang, Melissa H. Jia, Sathish Ponniah, Rodrigo Pedrozo, Camila Nicolli, Jeremy D. Edwards

## Abstract

One of the common mechanisms to trigger plant innate immunity is recognition of pathogen avirulence gene products directly by products of major resistance (*R*) genes in a gene for gene manner. In the USA, the *R* genes, *Pik-s, PiKh/m*, and *Pi-ta, Pi-39(t)*, and *Ptr* genes have been effectively deployed to prevent the infections of *M. oryzae* races, IB49, and IC17 for some time. *Pi-9* is only recently being deployed to provide overlapped and complimentary resistance to *Magnaporthe oryzae* races IB49, IC17 and IE1k in the USA. Pi-ta, Pi-39(t), Pi9 are major nuclear binding site-leucine rich (NLR) proteins, and Ptr is an atypical R protein with 4 armadillo repeats. AlphaFold is an artificial intelligence system that predicts a protein 3D structure from its amino acid sequence. Here we report genome sequence analyses of the effectors and avirulence (*AVR*) genes, *AVR-Pita* and *AVR-Pik*, and *AVR-Pi9*, in 3 differential *M. oryzae* races. Using AlphaFold 2 and 3 we find strong evidence of direct interactions of products of resistance genes *Pi-ta* and *Pik* with *M. oryzae* avirulence (*AVR*) genes, *AVR-Pita* and *AVR-Pik* respectively. We also found that AVR-Pita interacts with Pi-39(t) and Ptr, and Pi9 interacts with both AVR-Pi9 and AVR-Pik. Validation of direct interactions of two pairs of R and AVR proteins supported a direct interaction mechanism of plant innate immunity. Detecting interaction of both Ptr and Pi39(t) with AVR-Pita, and Pi-9 with both AVR-Pi9 and AVR-Pik, revealed a new insight into recognition of pathogen signaling molecules by these host R genes in triggering plant innate immunity.

## Introduction

Investigation of the molecular mechanisms of plant disease resistance is a long-term strategic plan to ensure that crops can be sustainably grown on the planet earth under changing conditions. Plants have evolved a sophisticated multifaceted mechanism to ward off biotic stress (Dangl and Jones, 2001). Upon the challenges by harmful microbes, plant resistance responses are initiated within minutes after pathogen invasion by resistance (*R*) genes (Zhou and Zhang, 2020). Plant *R* genes detect pathogens by a combination of indirect and direct recognitions in initiating a robust signaling cascade to inhibit pathogen growth and reduce damages of plant cells and organs. Rice blast disease is one of the most feared diseases for rice production worldwide. The causal agent *Magnaporthe oryzae* is one of the most genetically plastic pests and is highly adaptive to hosts (Dean et al. 2005). Rice plants, on the other hand, have evolved a complex innate immunity mediated by major *R* genes, and some of these *R* genes have been explored for crop protection. Several dozen *R* genes in rice have been cloned, and predicted proteins revealed that products of most *R* genes exhibit features of nucleotide binding site-leucine rich domain (NLR) (Wang et al. 2014).

The *R* genes in rice prevent the infections of *Magnaporthe oryzae* carrying the corresponding avirulence (*AVR*) genes (Silue et al. 1992). To date, more than 40 *M. oryzae AVR* genes have been mapped, 14 of which have been cloned. The cloned *AVR* genes are *PWL1* (Kang et al. 1995), *PWL2* (Sweigard et al. 1995), *AVR-Pita* (Orbach et al. 2000), *MoHTR1, MoHTR2* (Kim et al., 2020); *ACE1* (Böhnert et al. 2004), *AVR-Piz-t* (Li et al. 2009), *AVR-Pia* (Miki et al. 2009), *AVR-Pik/km/kp* (Yoshida et al. 2009), *AVR1-CO39* (Cesari et al. 2013), *AVR-Pii* (Fujisaki et al. 2015), *AVR-Pi9* (Wu et al. 2015a), *AVR-Pib* (Zhang et al. 2015) and *AVR-Pi54* (Ray et al. 2016). Among the cloned *AVR* and *R* gene pairs, *Pi-ta/AVR-Pita* (Jia et al. 2000; Orbach et al. 2000), *Pik/AVR-Pik* (Yoshida et al. 2009, Kanzaki et al. 2012), *Pia/AVR-Pia* (Miki et al. 2009; Ortiz et al. 2017), *Pi-CO39/AVR1-CO39* (Cesari et al. 2013) and *Pi54/AVR-Pi54* (Ray et al. 2016) can interact directly with each other. Other proteins, *Pii/AVR-Pii* (Fujisaki et al. 2015; Singh et al. 2016), *Piz-t/AVR-Piz-t* (Park et al. 2012; Park et al. 2016; Wang et al. 2016; Tang et al. 20170) were shown to indirectly detect pathogen *AVR* genes to induce disease resistance.

In the USA, despite the *R* gene *Pi-k* having been deployed since the 1960s, predominant rice varieties such as ‘Newbonnet’ were susceptible to the new races IB49 and IC17, resulting in several blast epidemics in the1980s. Blast *R* gene *Pi-ta* was introduced in the 1990s from a landrace variety ‘Tetep’ (Moldenhauer et al. 1990; Jia et al. 2019; McClung et al. 2020). Resistance specificity of *Pi-ta* to *M. oryzae* containing *AVR-Pita* has been well established: Firstly, the inheritance of blast resistance due to *Pi-ta* in the US rice varieties to the two major races, IB49 and IC17, was found (Jia et al. 2004). Later on, a wide range of additional global rice germplasm and weedy strains of rice were identified to carry the *Pi-ta* gene, and their functionality was verified with *M. oryzae* strains with and without *AVR-Pita* (Eizenga et al. 2006, Jia et al. 2004, Wang et al. 2008; Wang et al. 2004a,b; Wang et al. 2007; Wang et al. 2005; Lee et al. 2009; Lee et al. 2011; Li et al. 2014). In the 2000s, *M. oryzae* races IE1k found in commercial rice fields and IB33, a laboratory race, lacking *AVR-Pita* were infectious to *Pi-ta* containing rice varieties. In 2004, a rice variety ‘Banks’ was found to be susceptible in a commercial rice field. Blast isolates, B races and IE1k from Banks all have altered *AVR-Pita* alleles in virulent isolates, suggesting that resistance specificity of ‘Banks’ was abolished due to alteration of the *AVR-Pita* allele in a commercial field. ‘Banks’ was confirmed to carry the *Pi-ta* gene by *Pi-ta* specific markers (Jia et al. 2002) suggesting that *Pi-ta* mediated disease resistance was compromised by the new virulent B races (Zhou et al. 2007). The *AVR-Pita* allele from the original isolate 0-137 was introduced into two B races, B2 and B8, and the IE1k race resulted in resistance response to two US rice varieties, with *Pi-ta* verifying allele specificity of *AVR-Pita* to the *Pi-ta R* gene. The *Ptr* gene was named as *Pi-ta* required gene by finding a fast neutron induced susceptible mutant M2354 of Katy rice with *Pi-ta* to *M. oryzae* with *AVR-Pita* and was cloned using over 12K segregating progeny (Zhao et al. 1998). *Ptr* without *Pi-ta* confers blast resistance equivalent to that of *Pi-ta2* containing rice varieties. Rice varieties with *Pi-ta* were resistant to *M. oryzae* races IB49, IC17 with *AVR-Pita* and susceptible to *M. oryzae* race IE1k without *AVR-Pita* (Zhao et al. 2018). Deployment of *Pi9* was recently initiated after a gene specific DNA marker was developed (Scheuermann et al. 2019).

Pi-ta, Pik and Pi9 are NLR proteins, and Ptr is an atypical protein with 4 armadillo repeats located within 220 kb near the centromere of chromosome 12. Within *Pi-ta* and *Ptr* there is an additional *R* gene, *Pi39(t)* which is also a NLR protein (Liu et al. 2007). AlphaFold is a newly developed system to predict protein structure and their ability to interact with others based on amino acid sequences (Jumper et al. 2021; Abramson et al. 2024), and it can be used to verify protein-protein interactions and identify new interaction partners. The objectives of this study were to 1) analyze genomic structures of three differential blast races, IB49, IC17 and IE1k for their effectors, AVR genes, *AVR-Pita, AVR-Pik*, and *AVR-Pi9* and 2) examine pair wise protein and protein interactions to gain insight into mechanisms of initiating innate plant immunity.

## Materials and Method

### Fungus and growth

*Magnaporthe oryzae* races/isolates, IB49 (ML1), IC17 (ZN57), IE1k (TM2) were established on an oatmeal plate (Valent et al., 1986) with 10 sterilized filter discs from desiccated filter discs at - 20°C and grown in a fungal incubator at 26C, 37% relative humidity with continuous light (Percival, Perry, Iowa, USA) for 4-6 days until sporulated mycelia had grown on these discs. Filter discs with mycelia and spores of each race/isolate were sent to Mark Farman, University of Kentucky for genome sequencing. Genome sequence for IB49, IE1K and ICI7 were deposited at the NCBI genome section of *Pyricularia oryzae* (https://www.ncbi.nlm.nih.gov/datasets/genome/?taxon=318829). Genome assembly number for IB49 is ASM292504v1, for IE1K is ASM292498v1 and ASM292502v1 for IC17. While the genebank accession for IB49, IE1k and IC17 were CA_002925045.1, GCA_002924985.1 and GCA_002925025.1 respectively.

### Gene presence/absence analysis

We constructed a repeat library for each isolate using RepeatModeler version 2.0.5 (Flynn et al., 2020) and then soft masked three genomes with RepeatMasker version 4.1.5 (Smit et al., 2015). Genome annotation was performed using Braker3 version 3.0.8, which predicts genes based on both RNA data and a protein dataset (Gabriel et al. 2023). For RNAseq data, we used the datasets from NCBI BioProject PRJNA787662 and PRJEB45007, to include gene expression data across different life stages (Kim et al. 2022; Yan et al. 2023). Gene CDS and protein sequences were extracted using gffread version 0.12.7(Pertea et al. 2020). The completeness of genome annotation was evaluated with BUSCO version 5.4.5, with sordariomycetes_odb10 linkage dataset (Simao, et al. 2015). Subsequently, we grouped the pangenes of these three isolates using get_homologues-est version 3.6.2 with parameter “-t 0” to include all genes (Contreras-Moreira et al. 2017). Effector prediction was performed using effectorP version 3.0, with probability greater than 0.9 as positive (Sperschneider and Dodds 2022). To identify genes similar to known *AVR* genes *AVR-Pita, AVR-Pik* and *AVR-Pi9*, we used BLAST+ version 2.15.0 (Camacho et al. 2009) to align reference genes of *AVR-Pita* (Orbach et al. 2000), *AVR-Pik* (Yoshida et al. 2009) and *AVR-Pi9* (Wu et al. 2015a) to our representative pangenes.

### Receptor-ligand interactions prediction of with AlphaFold2 and AlphaFold3

We first tested the accuracy of AlphaFold2 (Evans et al. 2021; Jumper et al. 2021) and AlphaFold3 (Abramson et al. 2024) with nine biologically structured rice-related protein-protein interactions (PPIs) from the PDB database, which were not used in the training of AlphaFold models (Table 1). AlphaFold2-multimer version 2.3.2 was run using AlphaFold2 python wrapper AlphaPulldown version 1.0.4 (Yu et al. 2023) on Nvidia GPUs (A100 or V100). We further predicted the interactions of products of *AVR-Pita, AVR-Pik* and *AVR-Pi9* with products of rice resistance genes *Pi-ta* (Bryan et al. 2000), *Ptr* (Zhao et al. 2018), *Pik-p* (Yuan et al. 2011), *Pik(h)* (Sharma et al. 2005), *OsHIPP19* (Maidment et al. 2021) and *Pi9* (Qu et al. 2006) with AlphaFold2 and AlphaFold 3 (Jumper et al. 2021, Abramson et al. 2024). Preliminary screening was conducted with interface predicted template modelling score (ipTM) greater than 0.3 as candidate positive, and then we used AlphaPulldown to screen for likely true positive interactions, with thresholds ipTM greater than 0.6 and predicted template modelling score (pTM) greater than 0.5.

**Table 1.** Validation of AlphaFold2 and AlphaFold3 with untrained rice-related protein-protein interactions.

| PDB | AlphaFold2 |  | AlphaFold3 |  |
| --- | --- | --- | --- | --- |
|  | ipTM | pTM | ipTM | pTM |
| 7XPK | 0.94 | 0.93 | 0.89 | 0.91 |
| 7SA1 | 0.94 | 0.92 | 0.9 | 0.87 |
| 7XED | 0.87 | 0.90 | 0.75 | 0.83 |
| 7QPX | 0.85 | 0.86 | 0.83 | 0.83 |
| 7E40 | 0.81 | 0.69 | 0.74 | 0.63 |
| 8IF6 | 0.79 | 0.83 | 0.62 | 0.67 |
| 8B2R | 0.79 | 0.84 | 0.9 | 0.91 |
| 8R7A | 0.63 | 0.60 | 0.63 | 0.57 |
| 8R7D | 0.60 | 0.60 | 0.64 | 0.59 |

## Results

Braker3 annotation indicated that the isolate IB49 contains 10,648 genes, IC17 contains 10,619 genes and IE1K contains 10,608 genes, each with a BUSCO score of 99.1%, 99.2% and 99.2% respectively. A total of 11,015 non-redundant genes were identified across these three isolates (Table 2). Notably, 92% of the genes (10,089) are present in all three isolates, with only 429 genes shared by two isolates and 497 genes unique to one isolate: specifically, 287 in IB49, 117 in IC17 and 93 in IE1K (Table 2).

**Table 2.** Gene present and absent variations among three rice blast isolates.

| Category | Gene# | Ratio | Effector# | Ratio |
| --- | --- | --- | --- | --- |
| IB49 only | 287 | 3% | 9 | 3% |
| IC17 only | 117 | 1% | 14 | 5% |
| IE1K only | 93 | 1% | 5 | 2% |
| IB49 and IC17 only | 89 | 1% | 1 | 0% |
| IB49 and IE1K only | 105 | 1% | 3 | 1% |
| IC17 and IE1K only | 235 | 2% | 11 | 4% |
| IB49, IC17, and IE1K together | 10089 | 92% | 222 | 84% |
| Total | 11015 | 100% | 265 | 100% |

EffectorP predicted 265 effectors with a probability greater than 0.9 from non-redundant pangenes of these three isolates (Table 2). Of these, 84% of effectors were detected in all three isolates, with 28 unique effectors distributed as follows: 9 in IB49, 14 in IC17 and 5 in IE1K (Table 2). The effector AVR-Pita is shared between IB49 and IC17, AVR-Pik is present only in IB49 and AVR-Pi9 is found in all three isolates. The protein sequence of our representative AVR-Pik from IB49 is 100% identical with the AVR-Pik sequences reported by Martina Azelin and Dirum (2016). Additionally, the protein sequence of AVR-Pita from IB49 is 100% identical with AVR-Pita1, reported by Xing et al. (2013). The protein sequence of *AVR-Pi9* from IB49 is identical to the *AVR-Pi9* from reference genome 70-15 (Dean et al. 2005).

AlphaFold2 and AlphaFold3 successfully predicted all nine rice-related PPIs, achieving ipTM greater than 0.6 and pTM greater than 0.5 (Table 1). Structures predicted by AlphaFold models were very similar to these in the PDB database, especially for high-confidence PPIs like 7XPK (Figure 1A, 1B and 1C). For lower-confidence interactions, while most domain structures were similar to those documented in the PDB, some discrepancies were noted by AlphaFold models, like 8R7D, as shown in figure 1D, 1E and 1F. Among these nine pairs of PPIs, 8IF6 represents a complex of three protein interactions. Remarkably, the structures predicted by both AlphaFold2 and AlphaFold3 closely resemble the biological structures (Figure 1G, 1H, and 1I). Overall, AlphaFold2 and AlphaFold3 demonstrate comparable accuracy in predicting PPIs within the rice system.

**Figure 1.**
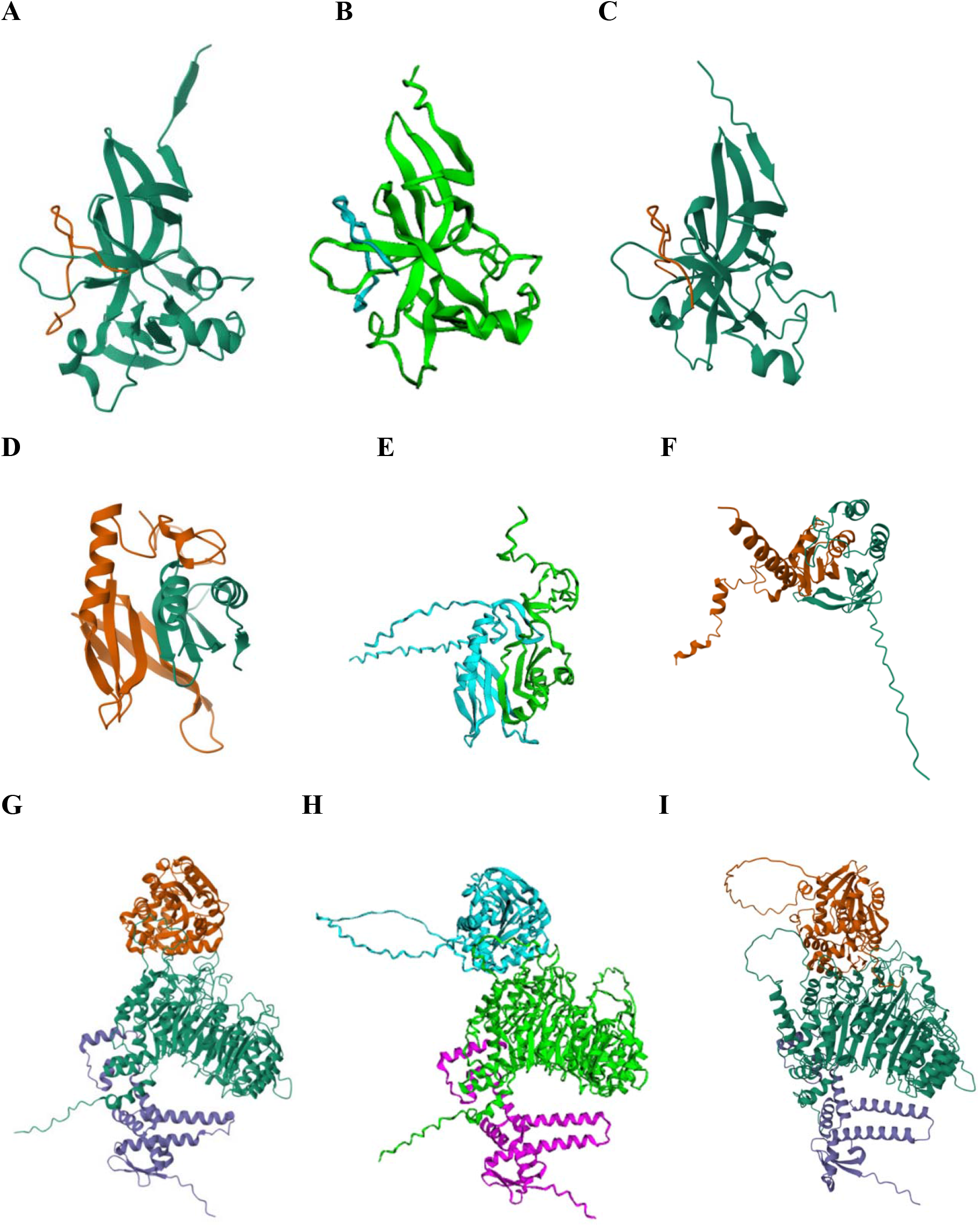
Comparative structures of rice-related protein-protein interactions as Documented in PDB and Predicted by AlphaFold models. (A) The PDB structures for 7XPK; (B) The predicted interaction structure of 7XPK by AlphaFold2; (C) The predicted interaction structure of 7XPK by AlphaFold3; (D) The PDB structures for 8R7D; (E) The predicted interaction structure of 8R7D by AlphaFold2; (F) The predicted interaction structure of 8R7D by AlphaFold3; (G) The PDB structures for 8IF6; (H) The predicted interaction structure of 8IF6 by AlphaFold2; (I) The predicted interaction structure of 8IF6 by AlphaFold3.

Furthermore, we predicted the interactions of products of *AVR-Pita, AVR-Pik* and *AVR-Pi9* with products of *R* genes, *Pi-ta, Ptr, Pik-p, Pik(h), OsHIPP19* and *Pi9* using AlphaFold2 and AlphaFold3. AlphaFold2 predicted the structures of interactions between products of *AVR-Pita* and products of *R* genes *Pi-ta* and *Pi39(t)* with both interactions primarily occurring through the leucine-rich repeat (LRR) domain, which spans amino acids 557 to 828 (Table 3 and Figure 2A, 2B and 2C). Additionally, AlphaFold2 provided insights into the interaction between products of *AVR-Pita* and *Ptr*, which involves an ARM repeat domain that is unique in its lack of an LRR domain (Figure 2D).

**Table 3.**
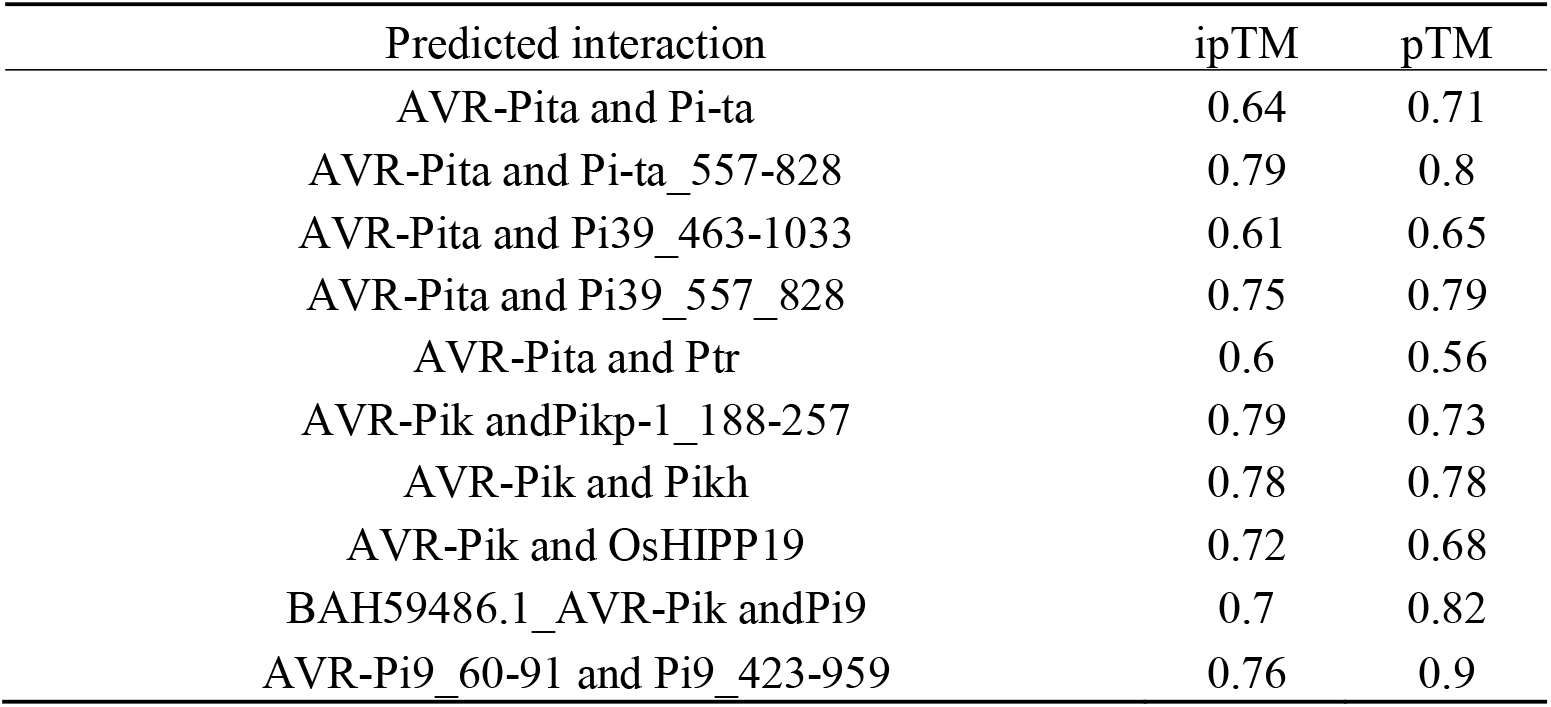
Predicted interactions statistics between AVR proteins and rice resistance genes by AlphaFold models.

**Figure 2.**
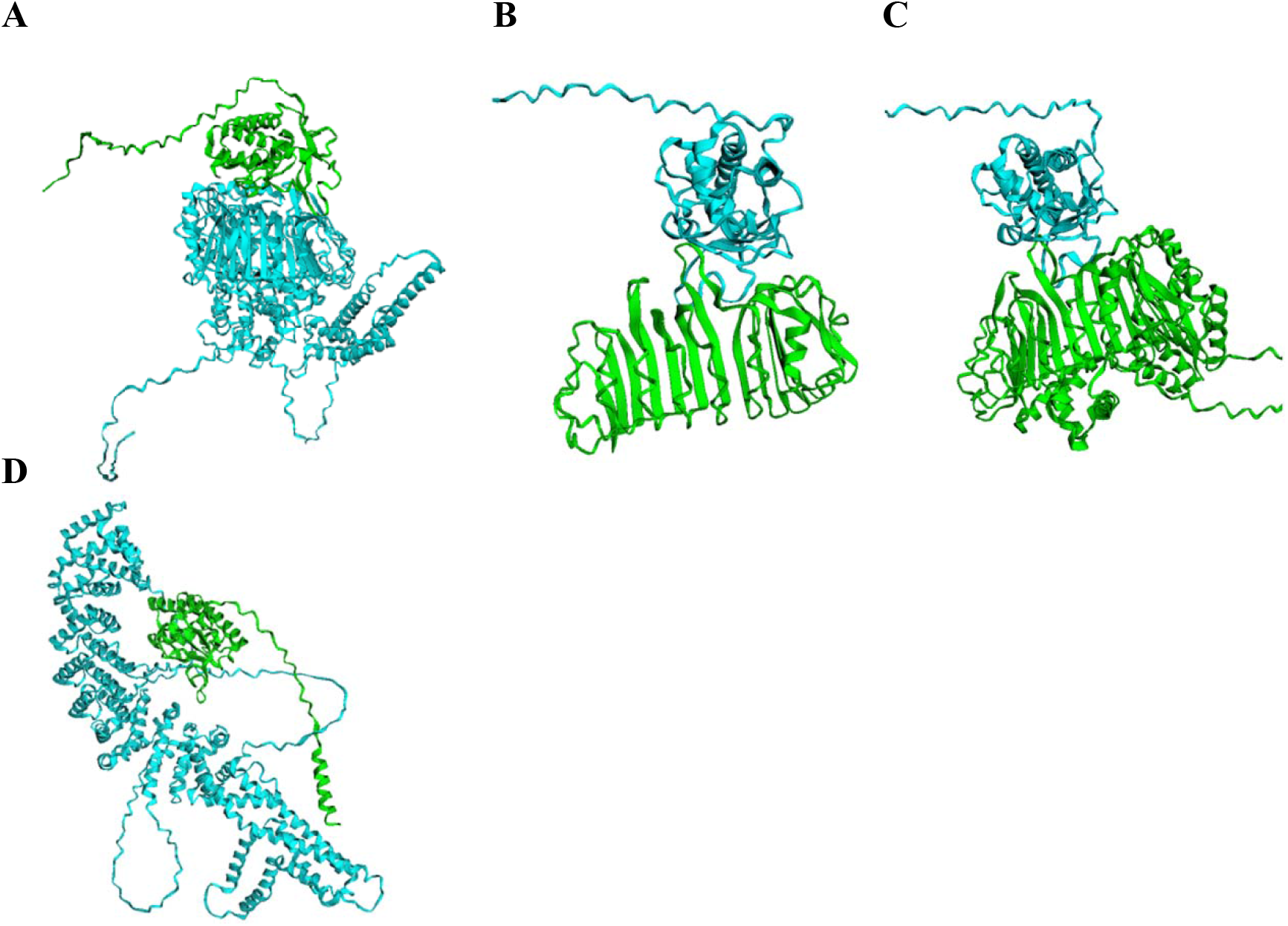
The interaction structures of AVR-Pita with three rice resistant genes Pi-ta, Pi39 and Ptr. (A) the interaction structure of AVR-Pita with Pi-ta; (B) Interaction structure of AVR-Pita with the leucine-rich repeat (LRR) domain of Pi-ta, specifically covering protein sequences from amino acids 557 to 828; (C) The interaction structure of AVR-Pita and Pi39; (D) The interaction structure of AVR-Pita with Ptr.

AlphaFold2 also successfully predicted the interaction of products of *AVR-Pik* with *Pik-p*,*OsHIPP19, Pik(h)*, and *Pi9* (Table 3 and Figure 3). Specifically, *AVR-Pik* recognizes Pik-p and OsHIPP19 through the HMA domain (Figure 3A and 3B). In contrast, both the Pik(h) and Pi9 proteins predominantly feature two continuous LRR domains with which AVR-Pik interacts (Figure 3C and 3D). Despite several attempts to model the interaction of AVR-Pik with Pi9, the best-case scenario yielded an ipTM of 0.56 and a pTM of 0.89, which narrowly missed our threshold of ipTM = 0.6 and pTM = 0.5. Subsequent attempts utilized a reference AVR-Pik protein sequence (GenBank accession number BAH59486.1, Yoshida et al. 2009), which differs by three amino acids from our AVR-Pik (Figure 4). These modifications enabled AlphaFold2 to predict the interaction between this reference AVR-Pik and Pi9, achieving a relatively high ipTM of 0.7 and pTM of 0.82 (Table 3 and Figure 3D). Meanwhile, AlphaFold3 effectively predicted the interaction structure between AVR-Pi9 and Pi9, as presented in Table 3 and Figure 5A and 5C. However, AlphaFold2 fell short in this endeavor, with the most favorable predictions reaching an ipTM of 0.56 and a pTM of 0.89, as shown in Figure 5B.

**Figure 3.**
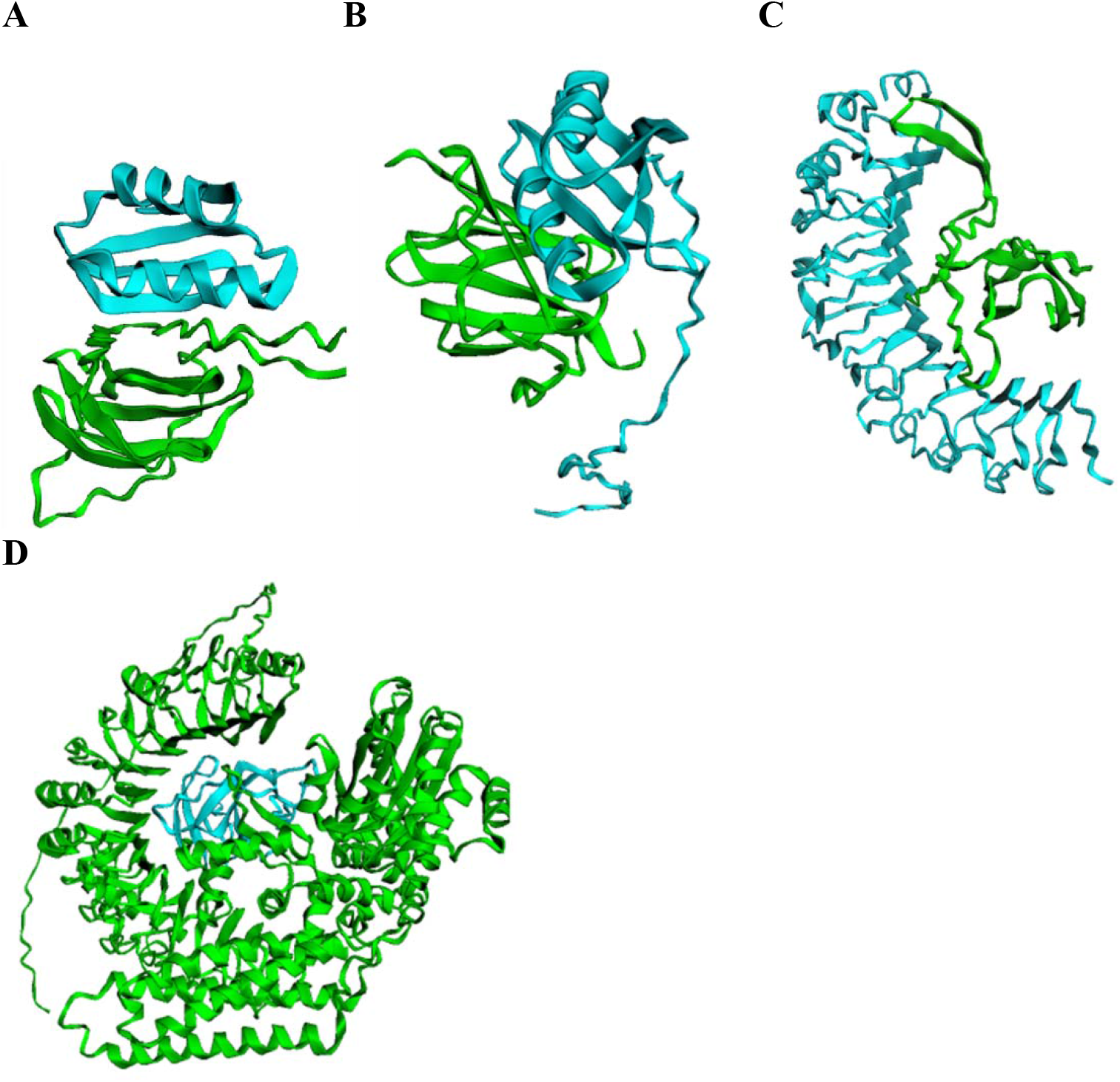
The interaction structure of AVR-Pik with four rice resistance genes Pik-p, OsHIPP19, Pik(h), and Pi9. (A) The interaction structure of AVR-Pik with the HMA domain of Pik-p; (B) The interaction structure of AVR-Pik with the OsHIPP19; (C) The interaction structure of AVR-Pik with Pik(h); (D) The interaction structure of AVR-Pik with Pi9.

**Figure 4.**
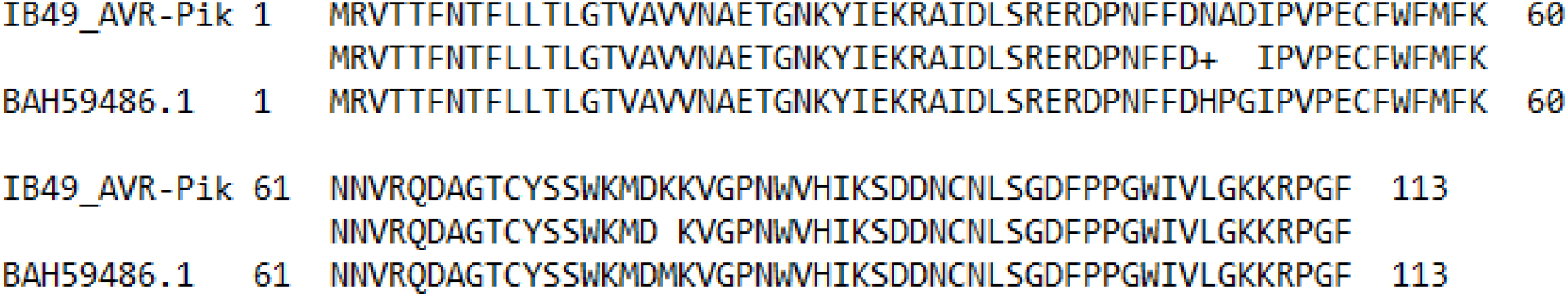
Amino Acid Sequence Comparison of Experimental and Reference AVR-Pik

**Figure 5.**
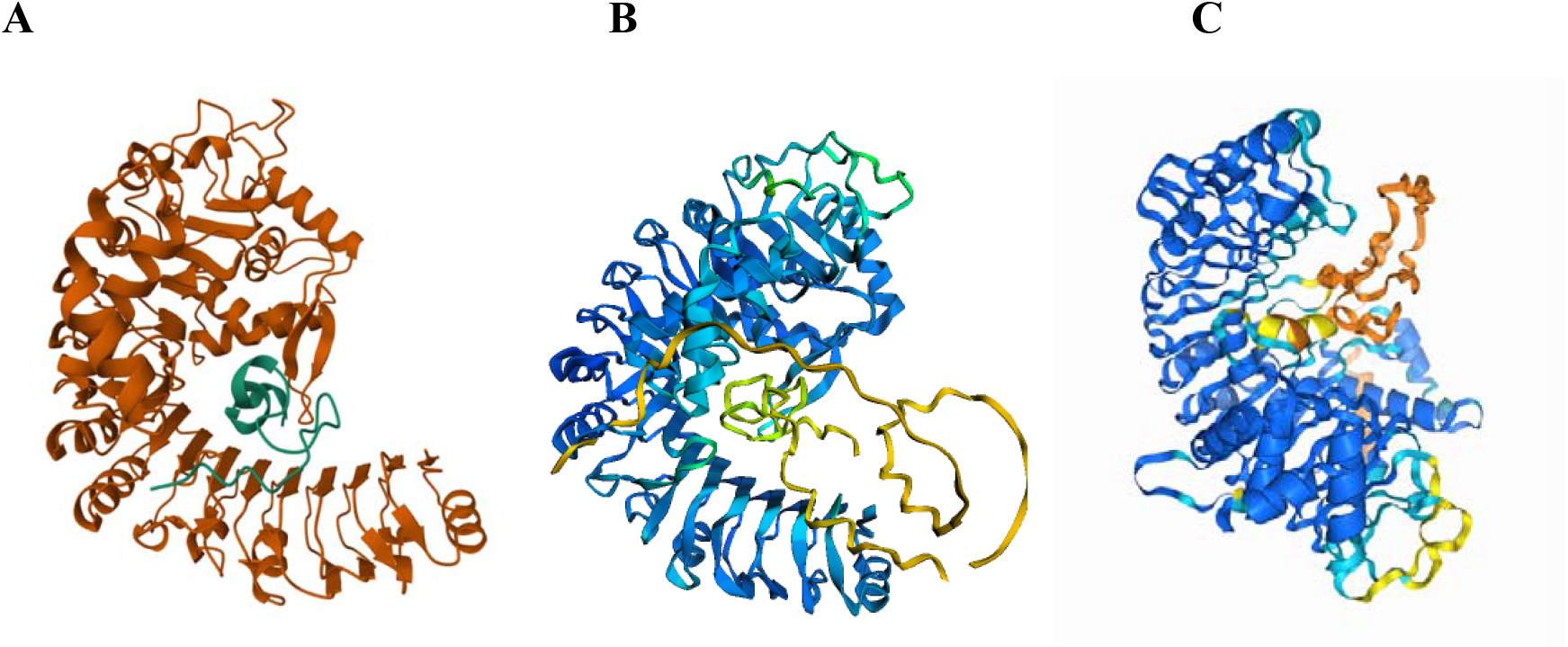
The interaction structure of AVR-Pi9 and Pi9 by AlphaFold2 and AlphaFold3. (A) Interaction structure of AVR-Pi9, covering protein sequences from amino acids 60 to 91, and Pi9, from amino acids 423 to 959, as predicted by AlphaFold3. The structure is color-coded to differentiate between the proteins; (B) Interaction structure of AVR-Pi9 and Pi9, spanning amino acids 423 to 959, as predicted by AlphaFold2. The structure is color-coded based on per-residue confidence scores (plDDT): dark blue indicates a plDDT greater than 90, signifying very high confidence; light blue represents a plDDT between 70 and 90, indicating confidence; yellow marks a plDDT between 50 and 70, suggesting low confidence; and orange signifies a plDDT below 50, indicating very low confidence in the structure of those residues; (C) Interaction structure of AVR-Pi9 and Pi9, covering amino acids 423 to 959, as predicted by AlphaFold3, using a similar plDDT color scale as in (B).

## Discussion

Historically, breeders have observed that rice varieties with *Pi-ta2* are resistant to more blast races than that of *Pi-ta*. All rice varieties with *Pi-ta2* also contain *Pi-ta*, suggesting that there are additional plant proteins involved in *Pi-ta2* mediated resistance. Rice varieties only with *Pi-ta*, such as Pi-no1, YT14 were resistant to *M. oryzae* races, IB49 and IC17 with *AVR-Pita*, susceptible to *M. oryzae* race IE1k without *AVR-Pita* (Zhao et al. 2018) suggesting that AVR-Pita is the pathogen signaling molecule for *Pi-ta* mediated disease resistance.

Since the release of AlphaFold2 in 2021, it has received over 23,000 citations and has been extensively utilized across various fields, including plant pathology. For instance, Homma et al. employed AlphaFold-Multimer to investigate the interactions between 1,879 small secreted proteins from seven tomato pathogens and six defense-related hydrolases of tomato. This study uncovered an effector hub targeting multiple microbial kingdoms. Additionally, Seong et al. explored the evolutionary mechanisms of effectors by predicting the structures of 26,653 effectors across 20 fungal and oomycete species using AlphaFold2 (Seong and Krasileva, 2023).

A comparative analysis of AlphaFold2 and AlphaFold3 highlights distinct advantages and limitations of these tools. AlphaFold2 offers greater versatility, primarily because its source code is publicly available, allowing for the customization of the MSA database and the adjustment of parameters such as the number of cycles and MSA depth, which influence prediction accuracy. In contrast, AlphaFold3, accessible only via a web service, restricts users to altering seed values. However, while AlphaFold2 requires substantial computational resources, AlphaFold3 can be accessed from standard computers and offers significantly faster processing times. Despite these differences, both versions successfully predicted the interactions of the nine tested PPIs.

Nonetheless, AlphaFold2 struggled with the low-quality structure prediction of AVR-Pi9 interacting with Pi9, whereas AlphaFold3 achieved acceptable predictive accuracy for the same interaction (Table 3 and Figure 5). NLR proteins are known to track ligand-products of *AVR* genes (Ngou et al., 2022). We reported the first detection of pathogen ligand AVR-Pita by a NLR protein, Pi-ta using in vitro and transient gene expression methods (Jia et al. 2000). AlphaFold revealed that AVR-Pita recognizes rice *R* proteins through two distinct mechanisms: The first involves recognition of the LRR domain, as evidenced in the interactions of AVR-Pita with two NLR proteins, Pi-ta and Pi39(t). The second mechanism involves pathogen recognition through the ARM domain, as seen in AVR-Pita interaction with Ptr (Table 3 and Figure 2). One plausible explanation is that AVR-Pita forms a bridge of a two NLR protein dimer and binds to the armadillo domain of Ptr protein in triggering resistance response. It is also possible that AVR-Pita is the ligand for Pi-ta, Pi39(t) and Ptr proteins complex in triggering effective resistance response. These results support that *Pi-ta* requires *Ptr* and *Ptr* is *R* gene independent to Pi-ta. Further validation will be required to establish domains/regions required for these interactions.

Similar to the AVR-Pita ligand, AVR-Pik recognizes proteins of *R* genes through two mechanisms. The HMA domain facilitates its interaction with Pik-p and OsHIPP19, a mechanism supported by multiple studies. De la Concepcion et al. (2018) first elucidated the biological structure of AVR-PikD and the HMA domains of Pikm-1 and Pikp-1, followed by Maidment et al. (2021), who described the structure of AVR-PikF and the HMA domain of OsHIPP19. Subsequently, Bialas et al. (2021) identified the structure of AVR-PikD with the HMA domain of Pik-1. These structures are preserved in the PDB database were likely used in training both AlphaFold2 and AlphaFold3, which explains the successful prediction of the interaction structure of AVR-Pik with Pik-p and OsHIPP19. In contrast, Pik(h) and Pi9, which lack an HMA domain but possess two continuous LRR domains, interact with AVR-Pik through these domains, as depicted in Figure 3C and 3D.

*Ptr* was identified from rice varieties and mutants with *Pi-ta* susceptible to *M. oryzae* containing *AVR-Pita* (Jia and Martin 2008) and was cloned using over 12k segregation progeny and validated using CRSPR-CAS9 (Zhao et al. 2018). Ptr is an atypical protein with 4 armadillo repeats lacking a U box, a feature resembling a typical E3 ligase. An E3 ligase APIP10 was known to directly interact with AVRPiz-t in transducing signals to a NLR receptor Piz-t in triggering resistance response (Park et al. 2016). Detecting of physical interaction of AVR-Pita with Ptr established a connection of indirect interaction of AVR protein, AVRPiz-t with R protein Piz-t. These findings revealed a remarkable feature of similarity and difference for each pair of *R* and AVR gene pair in triggering resistance response.

*Ptr* is a broad-spectrum *R* gene independent from *Pi-ta* and is required for *Pi-ta* in triggering effective resistance response; this suggests that *Ptr* is a failsafe for plant innate immunity (Zhao et al. 2018; Meng et al. 2020). The broad-spectrum *R* gene observed with *Ptr* may be associated with that of *Pi39(t)*, a gene encoding another NLR protein that is located within 12 kb from *Pi-ta* between *Pi-ta* and *Ptr*. All three *R* genes, *Pi-ta, Pi39(t)*, and *Ptr* have been bred into numerous resistant rice varieties in a linkage block globally (Jia 2009). The existence of a large linkage block at the *Pi-ta/Pi39(t)*/*Ptr* locus was one of the reasons that it took over 12K segregating progenies to clone *Ptr*. Similarly, Meng et al. (2020) suggested that *Ptr* is allelic to *Pi-ta2*, and Greenwood et al. (2024) showed resistance to some strains of *M. oryzae* in some global resistance varieties was due to *Ptr*. Further fine mapping and functional validation each one and in combination of them will be required to determine if *Ptr* is an allelic to *Pi-ta2*.

Genes involved in resistance to pathogens are often found in a cluster within a short physical distance. *Pi-ta, Pi39(t)* and *Ptr*, all within a 220 kb region on chromosome 12 of rice, are responsible for blast resistance. Using AlphaFold we showed that one NLR protein and one armadillo repeat-containing protein are involved in detecting pathogen signal molecule AVR-Pita in triggering effective disease resistance response. Further validations and elucidation of interaction domains would pave the way for precise engineering of resistance to blast fungus, a notable cereal killer.

## Acknowledgements

The authors are grateful for the technical support of Heather Box, Paul Braithwaite and other staff members of DB NRRC. For DNA sequencing we thank Dr. Mark Farman (University of Kentucky, Lexington). This work is in part supported by Nation Science Foundation, Grant Award Number: IOS-1947609 and USDA SCINet Postdoctoral fellowship. This research used resources provided by the SCINet project and the AI Center of Excellence of the USDA Agricultural Research Service, ARS project numbers 0201-88888-003-000D and 0201-88888-002-000D. USDA is an equal opportunity provider and employer.

## Author contributions

YJ, JE, LW, AO, and YH designed the study. LW, AO, YH, MHJ, RP, CN performed research. LW analyzed the data, LW and YJ wrote the manuscript. All authors read and approved the final submission.

## CONFLICT OF INTEREST STATEMENT

The authors declare no conflict of interest.

## DATA AVAILABILITY STATEMENT

Genome sequences of blast races are available at https://www.ncbi.nlm.nih.gov/datasets/genome/?taxon=318829. Genome assembly number for IB49 is ASM292504v1, for IE1K is ASM292498v1 and ASM292502v1 for IC17. While the genebank accession for IB49, IE1k and IC17 were CA_002925045.1, GCA_002924985.1 and GCA_002925025.1 respectively.

## Notes

### Competing Interest Statement

The authors have declared no competing interest.

